# Navigating the phenotype frontier: The Monarch Initiative

**DOI:** 10.1101/059204

**Authors:** Julie A. McMurry, Sebastian Köhler, James P. Balhoff, Charles Borromeo, Matthew Brush, Seth Carbon, Tom Conlin, Nathan Dunn, Mark Engelstad, Erin Foster, JP Gourdine, Julius O.B. Jacobsen, Dan Keith, Bryan Laraway, Suzanna E. Lewis, Jeremy Nguyen Xuan, Kent Shefchek, Nicole Vasilevsky, Zhou Yuan, Nicole Washington, Harry Hochheiser, Christopher J. Mungall, Tudor Groza, Damian Smedley, Peter N. Robinson, Melissa A. Haendel

## Introduction

The principles of genetics apply across the entire tree of life. At the cellular level we share biological mechanisms with species from which we diverged millions, even billions of years ago. We can exploit this common ancestry to learn about health and disease, by analyzing DNA and protein sequences, but also through the observable outcomes of genetic differences, i.e. phenotypes.

To solve challenging disease problems we need to unify the heterogeneous data that relates genomics to disease traits. Most databases tend to focus either on a single data type across species, or on a single species across data types. Although each database may provide rich, high-quality information, none is unified and comprehensive across species, over biological scales, and throughout data types (Figure 1A).

**Figure 1.**
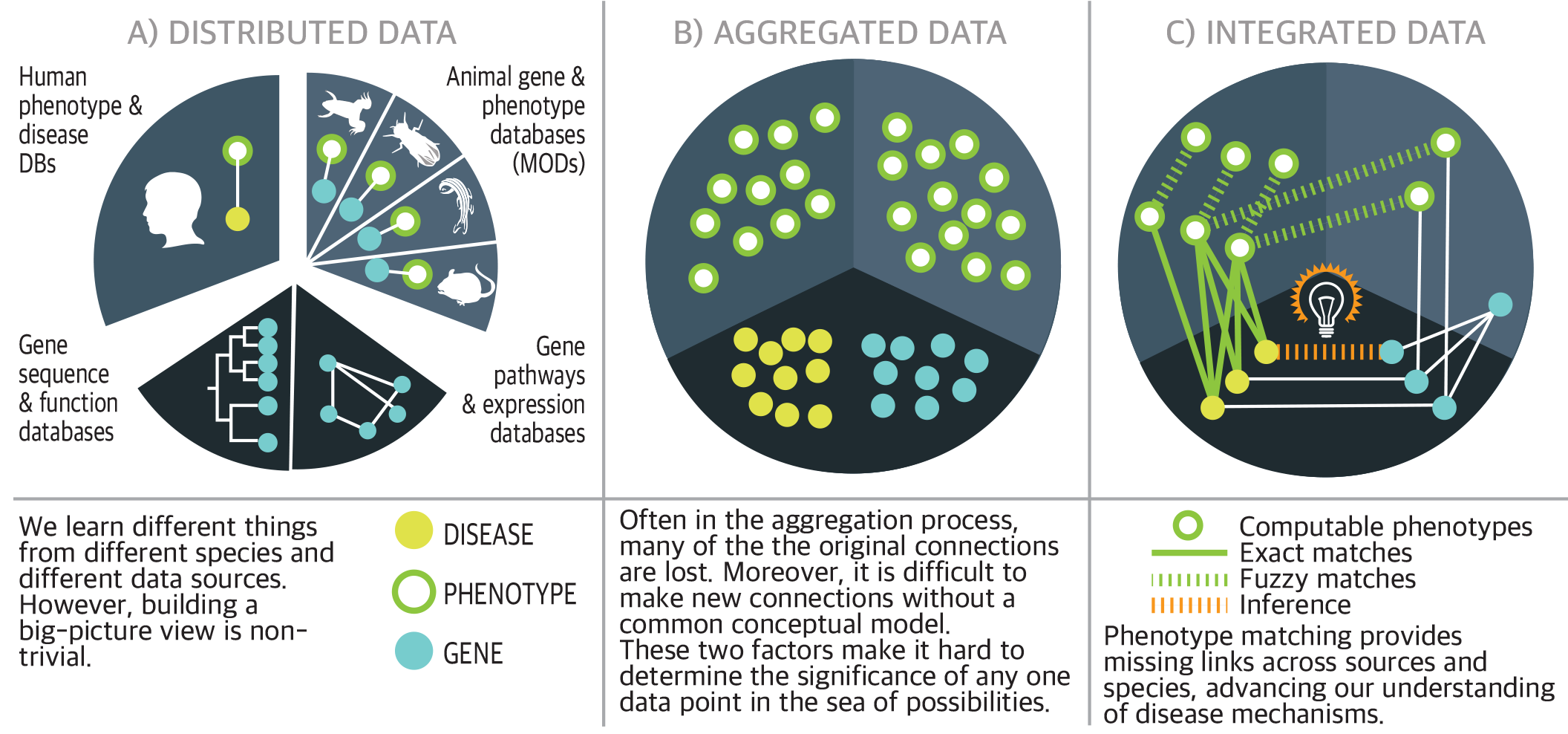
Role of phenotypes in data integration.

Without a big-picture view of phenotypic data, many questions in genetics are difficult or impossible to answer. Use of computable phenotypes—which can be efficiently analyzed with algorithms—is a crucial strategy for gaining this broader view. When a disease has an unknown genetic basis, or is associated with mutations in multiple genes, computable phenotypes can provide valuable clues to the underlying complexity. Aggregating the data in one place is necessary for search and retrieval (Figure 1B), but aggregation often results in a loss of data richness and meaning. *Connecting* the dots enables the bigger picture to emerge; computable phenotypes (Figure 1C) provide key links across sources, species, and data types.

The Monarch Initiative (https://monarchinitiative.org) provides tools for genotype-phenotype analysis, genomic diagnostics, and precision medicine across broad areas of disease. These tools depend on the data integrated through computable phenotypes for cross species comparisons.

To fully exploit the power of computable phenotypes, several obstacles must be overcome. These are illustrated for a specific example related to a family of diseases in Figure 2.

**Figure 2.**
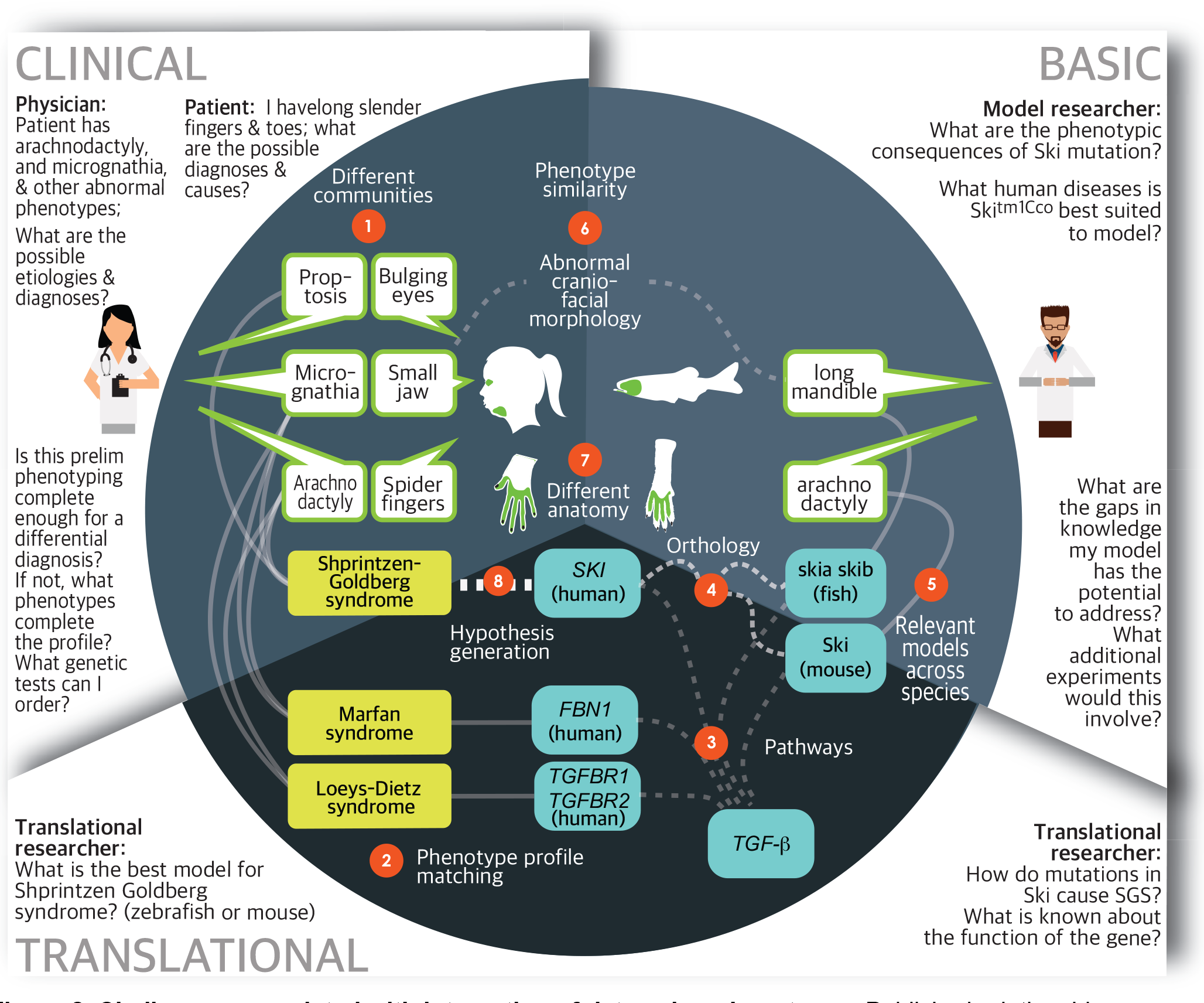
Challenges associated with integration of data using phenotypes. Published relationships are shown in solid lines. Dashed lines show relationships that require computation and/or data integration. Around the perimeter of the figure are examples of the types of questions that are difficult to answer using traditional (non-integrative) methods. These questions are divided into “clinical”, “basic”, and “translational” research categories. Each challenge is explained further below.

1. **Different communities:** Different communities use different language to describe the same phenotypes, even for the same species (e.g. clinicians use the term ‘Micrognathia,’ where patients would use ‘small jaw’). Similarly, a mouse researcher might describe the orthologous phenotype as ‘small mandible’.
2. **Phenotype profile matching:** Many diseases closely resemble each other, and the constellation of phenotypes associated with any given disease rarely, if ever, manifest in the same way in every affected patient. Identifying the hallmark features as well as key differentiating phenotypes is an essential part of differential diagnosis. In the above example, ‘proptosis’ is common in Shprintzen Goldberg syndrome, but not in Loeys-Dietz or Marfan syndromes. Any observed phenotype profile varies depending on the location and nature of the gene variation. This degree of variability means that fuzzy matching between sets of phenotypes can play a key role, both in differential diagnosis and mechanistic inquiry.
5. **Relevance of pathways and co-involvement:** Knowing which genes/proteins are co-implicated in similar phenotypes provides important clues. Some evidence in mice shows that mutations in *FBN1* as well as *TGFBR1* and *TGFBR2* lead to an overlapping set of abnormal phenotypes including those of the skeletal system (Arslan-Kirchner *et al.* 2016). Alteration of the murine ortholog of *SKI* (Ski) is associated with skeletal craniofacial anomalies in mouse models, and the *SKI* gene plays a role in *TGF* metabolism.
6. **Orthology:** Gene orthology between species is not black and white: some sequences are more closely related than others; moreover, describing orthology is complicated by factors such as gene duplication and splice variation. Sources differ as to whether given genes are orthologs and if so, what type (e.g. least diverged ortholog, paralog, 1-to-1, many-to-many, etc.) (O'Brien *et al.* 2005, Altenhoff *et al.* 2016). In the above example, the zebrafish has two copies of the *SKI* ortholog, skia and skib.
7. **Relevant models across species:** A single animal model rarely recapitulates all of the phenotypes exhibited in human disease. It often takes a combination of models to help form a complete picture. In this case, the Marfan mouse model exhibits the ‘arachnodactyly’ whereas the zebrafish exhibits the craniofacial abnormalities (Doyle *et al.* 2012).
8. **Atomic phenotype similarity:** Unlike genes, phenotypes are not discrete entities; this makes querying databases for phenotypes a difficult problem related to granularity. For instance, queries for any term (e.g. ‘hyperkeratosis’) should contain all results associated with more specific variations of the underlying concept (e.g. ‘palmoplantar hyperkeratosis’). It is not only hierarchical relationships that matter, but basic similarity. For instance, ‘micrognathia’ is similar to ‘small mandible’ but neither of these is a parent of the other; rather both terms descend from ‘abnormal jaw morphology’.
9. **Anatomy, and biological scales:** Similar phenotypes are recorded in different species for analogous anatomical regions (e.g. ‘hand’ versus ‘paw’). They also apply to different scales (e.g. ‘neurological phenotype’ is related to more specific concepts such as ‘dopaminergic cell loss’). Structuring these concepts into networks allows both machines and humans to navigate complex interlinked data.
10. **Hypothesis generation:** Simultaneously overcoming challenges 1-7 enables us to generate new hypotheses. For instance, we could speculate that mutation in the human *SKI* gene might lead to a disease with similarities to Marfan syndrome and Loeys-Dietz syndrome. In fact, these considerations supported the discovery of mutations in the *SKI* gene as the cause of Shprintzen Goldberg syndrome (Schepers *et al.* 2015).

Additional biological complexities make it even harder to build a complete and accurate picture with the available phenotypic information. To name a few not illustrated above:

11. **Inference:** Phenotypes are often associated with diseases and diseases to genes; thus the relationship between a specific phenotype and a specific gene may need to be inferred. For instance, if we know that *FBN1* is implicated in Marfan syndrome and that skeletal anomalies such as ‘arachnodactyly’ are associated with Marfan syndrome, then we can infer that *FBN1* is likely to play some role in skeletal development and homeostasis.
12. **Staging, severity:** Interpretations are affected by the stage of an organism or the stage of disease at which the phenotype is observed, in combination with phenotypic severity.
13. **Time:** Biological processes occurring at different developmental times are hard to compare across organisms.
14. **Phylogenetic distance:** A model organism may not present the exact spectrum of phenotypes when faced with an orthologous genetic variation, and the similarities between phenotypes become subtler and thus harder to find and quantify as phylogenetic distances increases (e.g. pleiotropic phenotypes in Bardet-Biedl are similar to effects seen in the cilia and basal bodies of single-celled eukaryotes).
15. **Noise:** Observations may be incomplete or artefactual (noise).

**A common conceptual framework.** Data scientists often apply ontologies to organize heterogeneous data. Ontologies are collections of concepts logically organized and linked. Most anatomy, phenotype, and disease ontologies describe the biology of one particular species. Examples are the Human Phenotype Ontology (HPO) (Kohler *et al.* 2014) and the Mouse Anatomy Ontology (Hayamizu *et al.* 2015). Monarch has developed four species-agnostic ontologies designed to unify their species-specific counterparts: GENO for genotypes (Brush *et al.* 2013), Uberpheno for phenotypes (Kohler *et al.* 2013), UBERON for anatomy (Haendel *et al.* 2014), and MONDO for diseases (Mungall *et al.* 2016). These ontologies provide a bridge between species/domain-specific ontologies, allowing unified analysis of disparate data sources (Figure 3). And Monarch contributes to the Gene Ontology, which also unifies gene function and subcellular anatomy across species (Ashburner *et al.* 2000).

**Figure 3.**
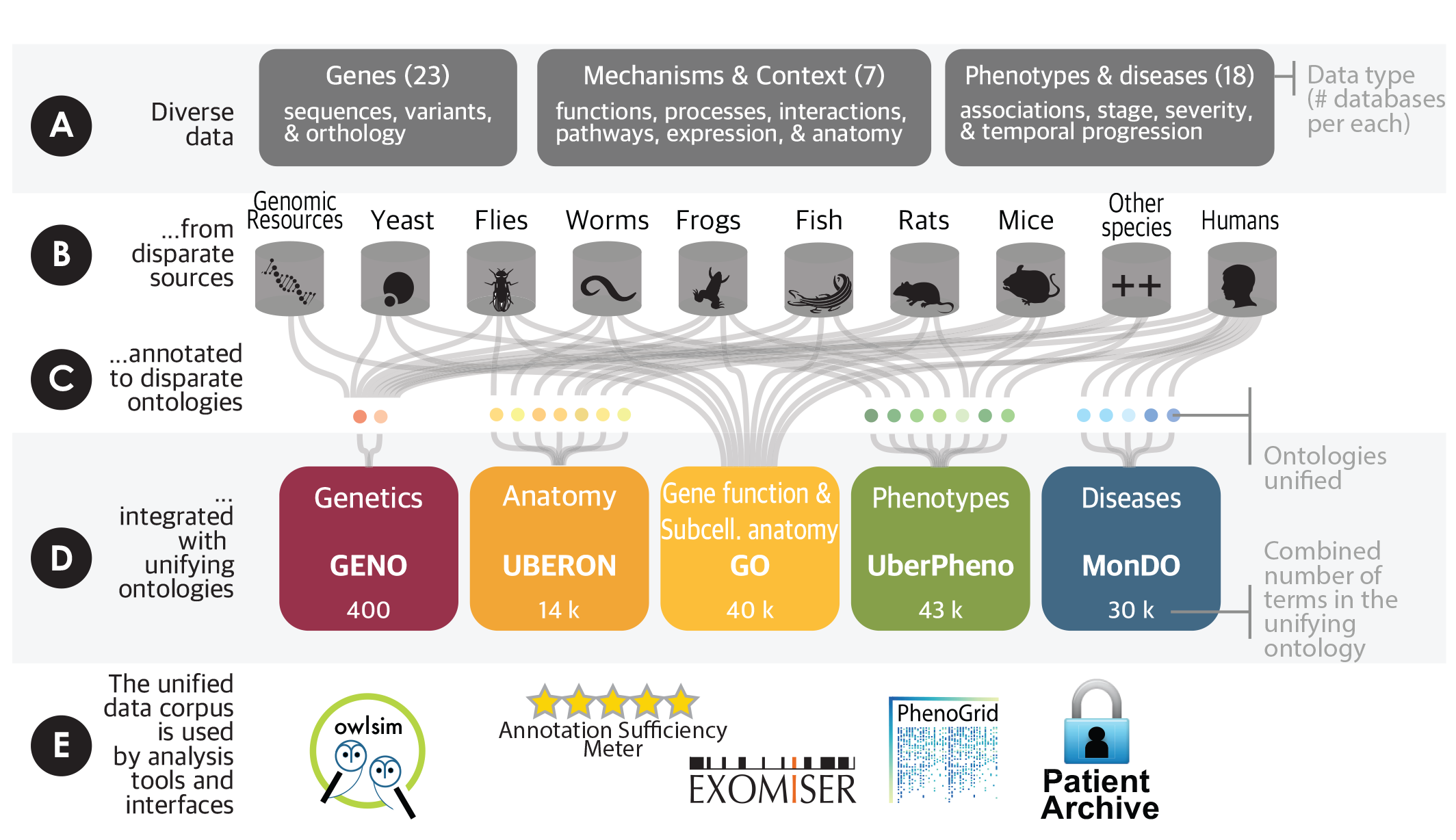
Monarch's ontology-driven data integration pipeline. Diverse data from disparate sources and annotated to disparate species-specific ontologies is integrated with unifying ontologies. The unified data corpus is used by analysis tools and interfaces.

Monarch tools leverage this conceptual framework to help users understand and diagnose disease. Statistical similarity calculations enable comparison across species (Fig 2.7), biological scales (Fig 2.9), and community-specific vocabularies (Fig 2.1) (Smedley *et al.* 2013). Monarch supports researchers and clinicians using this data with visualization tools, Application Programming Interfaces (APIs), and a rich web site (<u>https://monarchinitiative.org</u>). These approaches make it possible to overcome limitations in the data for many applications, including disease diagnostics (Bone *et al.* 2015), drug repurposing, and improved phenotyping, both clinically and in model organisms (e.g. helping identify candidate phenotyping assays based on preliminary phenotyping). Indeed, Monarch's unified data corpus and tools have been applied to diagnosing real patients, and plans are underway to scale up their use with larger efforts, including the Undiagnosed Diseases Network (UDN) (Brownstein *et al.* 2015) and the 100,000 Genomes Project (http://www.genomicsengland.co.uk/the-100000-genomes-project/).

To achieve this vision, we need technological advances and collaborative processes beyond the common conceptual framework. Existing descriptions of phenotypes and their relationships to genomic variations are all-too-frequently provided in community-specific formats, which lack the details and computational meaning needed for integration. Although we have made some progress with natural language approaches to extracting key details (Groza *et al.* 2015), expensive manual curation is still necessary.

To increase the portability and computability of phenotype descriptions, data providers and journals should use common phenotype information models. Such models require proper identification of the organisms being phenotyped. We have shown that approximately 33% of mouse strains and 13% of fish strains were not uniquely identifiable in the literature, causing the associated phenotype data to be lost to public repositories (Vasilevsky *et al.* 2013). Increased use of organism-specific nomenclature and identifiers, as supported by the Model Organism Databases, will be necessary for more effective sharing of phenotype data.

A standard data exchange format is needed to ensure that phenotypic knowledge is computable and accessible across a variety of sources. Toward this end, we are developing an exchange format (<u>phenopackets.org</u>) that will do for phenotype data what existing formats (e.g. FASTA, VCF, BED) have done for sequence data. Phenopackets can be used in a variety of settings, such as for submission to journals, in public databases, for biodiversity collections, and for clinical data sharing. They can apply to one organism or to groups of organisms, and for qualitative or quantitative data.

We invite the community to aid in the sharing, aggregation, and integration of cross-species phenotype data. By using, testing, and contributing to the phenopacket standard, you will be connecting the very dots that maximize mechanistic discovery of the genetic bases of health and disease.

## References

Arslan-Kirchner, M., E. Arbustini, C. Boileau, P. Charron, A. H. Child et al., 2016 Clinical utility gene card for: Hereditary thoracic aortic aneurysm and dissection including next-generation sequencing-based approaches. Eur J Hum Genet 24: e1–5.

Ashburner, M., C. A. Ball, J. A. Blake, D. Botstein, H. Butler et al., 2000 Gene ontology: tool for the unification of biology. The Gene Ontology Consortium. Nat Genet 25: 25–29.

Bone, W.P., N. L. Washington, O. J. Buske, D. R. Adams, J. Davis et al., 2015 Computational evaluation of exome sequence data using human and model organism phenotypes improves diagnostic efficiency. Genet Med.

Brownstein, C. A., I. A. Holm, R. Ramoni and D. B. Goldstein, 2015 Data sharing in the undiagnosed diseases network. Hum Mutat 36: 985–988.

Brush M. H., C. J. Mungall, N. Washington, and M. A. Haendel, 2013 What's in a Genotype?: An Ontological Characterization for Integration of Genetic Variation Data. International Conference on Biological Ontology.

Doyle, A. J., J. J. Doyle, S. L. Bessling,S. Maragh, M. E. Lindsay et al., 2012 Mutations in the TGF-beta repressor SKI cause Shprintzen-Goldberg syndrome with aortic aneurysm. Nat Genet 44: 12491254.

Groza, T., S. Kohler, D. Moldenhauer, N. Vasilevsky, G. Baynam et al., 2015 The Human Phenotype Ontology: Semantic Unification of Common and Rare Disease. Am J Hum Genet 97: 111–124.

Haendel, M. A., J. P. Balhoff, F. B. Bastian, D. C. Blackburn, J. A. Blake et al., 2014 Unification of multispecies vertebrate anatomy ontologies for comparative biology in Uberon. J Biomed Semantics 5: 21.

Hayamizu, T. F., R. A. Baldock and M. Ringwald, 2015 Mouse anatomy ontologies: enhancements and tools for exploring and integrating biomedical data. Mamm Genome 26: 422–430.

Kohler, S., S. C. Doelken, C. J. Mungall, S Bauer, H. V. Firth et al., 2014 The Human Phenotype Ontology project: linking molecular biology and disease through phenotype data. Nucleic Acids Res 42: D966–974.

Kohler, S., S. C. Doelken, B. J. Ruef, S Bauer, N Washington et al., 2013 Construction and accessibility of a cross-species phenotype ontology along with gene annotations for biomedical research. F1000Res 2: 30.

Mungall, C. J., S Koehler, P Robinson, I Holmes, and M. A. Haendel, 2016 k-BOOM: A Bayesian approach to ontology structure inference, with applications in disease ontology construction. bioRxiv.

O'Brien, K. P., M Remm and E. L. Sonnhammer, 2005 Inparanoid: a comprehensive database of eukaryotic orthologs. Nucleic Acids Res 33: D476–480.

Schepers, D., A. J. Doyle, G. Oswald, E. Sparks, L. Myers et al., 2015 The SMAD-binding domain of SKI: a hotspot for de novo mutations causing Shprintzen-Goldberg syndrome. Eur J Hum Genet 23: 224–228.

Smedley, D., A Oellrich, S Kohler, B Ruef, M Westerfield et al., 2013 PhenoDigm: analyzing curated annotations to associate animal models with human diseases. Database (Oxford) 2013: bat025.

Vasilevsky, N. A., M. H. Brush, H. Paddock, L. Ponting, S. J. Tripathy et al., 2013 On the reproducibility of science: unique identification of research resources in the biomedical literature. PeerJ 1: e148.

